# Protamine sequence determines species-specific nuclear shape and histone retention

**DOI:** 10.1101/2025.03.20.644273

**Authors:** Marta Czernik, Luca Palazzese, Dawid Winiarczyk, Domenico Iuso, Saadi Khochbin, Josef Fulka, Helena Fulka, Rocío Villafranca, Juan Antonio Rielo, Ana Sanchez-Rodriguez, Nerea Latorre, Clara Agudo-Rios, Eduardo R. S. Roldan, Maria Eugenia Teves, Pasqualino Loi

**Author notes:** equal controbuttion. Corresponding Authors: Eduardo Roldan, Department of Biodiversity and Evolutionary Biology, Museo Nacional de Ciencias Naturales (CSIC), Madrid, Spain, Pasqualino Loi, Laboratory of Embryology, Department of Veterinary Medicine, University of Teramo, Campus Sant’Agostino, Via Balzarini 1, 64100, Teramo, Italy.

## Abstract

Nuclear shape observed after the forced expression of mouse or human Protamine 1 (PRM1) in fibroblasts led us to propose the hypothesis that PRM1 sequence plays an important role in imposing the overall shape of the protaminized nucleus. Comparison of mouse and human PRM1 sequence pointed to cysteines 15 and 29 as potential critical residues in the mouse PRM1 sequence inducing the characteristic mouse “hooked” nuclear sperm shape. To explore this idea, mice with mutations in PRM1 Cys15 and Cys29 were generated. These mice remained fertile with no significant changes in sperm count or protamine expression levels. However, modifications in sperm head shape were observed. Transmission electron microscopy revealed disrupted chromatin condensation in mutant sperm, with several morphological changes and a remarkable increase in histone retention. Overall, the findings suggest that species-specific PRM1 cysteine residue positions are crucial for nuclear shape determination and histone retention in spermatozoa.

## Introduction

In contrast to the invariable spherical shape of the oocyte, the morphology of spermatozoa varies widely across species, particularly with regards to head shape (Roldan et al. 1992, Breed et al. 2014, Teves and Roldan 2022). Several selective forces have been proposed to account for sperm diversity, including post-copulatory sexual selection in the forms of sperm competition or cryptic female choice (Roldan et al. 1992, Teves and Roldan 2022). Thus, when spermatozoa from rival males compete to gain fertilizations, changes in sperm head shapes may confer hydrodynamic advantages in swimming (Tourmente et al. 2011, Gómez Montoto et al. 2011) or in sperm interactions with the female tract (Varea-Sánchez et al. 2016). Selection by females may also be based on sperm morphology and, therefore, some head morphs may become better suited to negotiate barriers in the female tract (Firman et al. 2016).

Studies in the mouse have demonstrated that the cytoskeleton plays a pivotal role in shaping nuclear morphology during sperm differentiation. At the beginning of the spermatid elongation phase the nucleus and the early acrosome polarize to one end of the cell (O’Donnell, 2015). As early as in step 5 of spermiogenesis, the F-actin filaments and associated proteins forming the acroplaxome prime nuclear shaping. Next, tensions generated by the microtubules and actin filaments in the manchette and the interaction with the nuclear envelope through the LInker of Nucleoskeleton and Cytoskeleton (LINC) complex provide a major shaping factor (Fayomi and Orwig, 2018). At this point, the nucleus compacts progressively, as the nucleosomal chromatin is transformed into compacted fibres by the sequential replacement of histones by testis-specific histone variants, transition proteins (TPs) and protamines (PRMs) (either protamine 1 alone in most mammals, or protamines 1 and 2 in primates and rodents and a few other species, Barral et al. 2017).

However, the possibility that chromatin compaction could have a more relevant role in nuclear shaping has been already suggested (Teves and Roldan, 2022). Our previous studies exploring strategies for nuclear reprogramming of somatic cell nuclei (Loi et al. 2016) have found that, when somatic cells express a heterologous mouse or human protamine 1 gene (*Prm1, PRM1*), their nuclei acquire a sperm-like shape reminiscent of the species; that is, curved in fibroblasts expressing mouse *Prm1*, and straight with human *PRM1* (Iuso et al. 2015, Czernik et al. 2016, Palazzese et al. 2018). These findings suggest that protamine sequence might play a role in shaping sperm nuclei. The PRM1 protein sequence is not well conserved between mammals, which can lead to different 3D structure of the protein in various species. Mammalian PRM1 is typically 50 amino acids long and it is made of a central arginine-rich domain that binds to 11 bp located in the major groove of the DNA (Balhorn, 1991). Both ends of PRM1 have a cysteine-rich domain. Intra- and inter-protamine disulfide bonds between cysteine residues are responsible for the condensation of DNA in the final maturation phases of the male gamete. However, how the DNA-bound protamines cross-link through S-S bonds and compact remains a matter of debate. Moreover, the surprising lack of crystallographic data of protamine/DNA complexes further complicates modelling work.

In this study, we hypothesized that the position of these S-S bonds along the protein might have an impact on the overall structure and shape of the resulting compacted genome. To test this hypothesis, we first explored whether there are differences in nuclear remodeling depending on the source of PRM1 (mouse, *mPrm1*, or human, *hPRM1*) by quantifying changes in nuclear shape induced by protamine expression in mouse or sheep fibroblasts. Because mouse and human PRM1 differ in the number and position of cysteine residues, we also examined if presence/absence of certain cysteines affect nuclear remodeling. To this end, we progressively substituted cysteine residues in the mouse gene by site-directed mutagenesis and expressed the mutant vectors in sheep and mouse fibroblasts. Based on these findings, PRM1 mutant mice were generated by CRISPR and used to investigate the effect of the lack of cysteine residues in the mouse protamine 1 sequence on sperm head morphological traits, nuclear compaction and fertility. Our findings suggest that, while not compromising overall fertility, mouse PRM1 mutant spermatozoa display head morphological differences, in shape as well as in thickness, probably as consequence of a change in nuclear compaction. A reduced compaction might have been the consequence of a higher histone retention in mutated spermatozoa, in comparison to the control. Overall, our data indicate that protamine sequence affects head morphology and the protamine to histone ratio in spermatozoa, and raise the possibility that protamine sequence, specifically the number and the position of cysteines, might be important for the wide morphological variations of sperm heads across species.

## Results

### Protamine 1 Drives Fibroblast Nuclear Remodeling and the Role of Cysteine Residues in Nuclear Shape

We have shown that the expression of protamine 1 (PRM1) significantly influences nuclear morphology in fibroblasts, reflecting species-specific sperm nuclear shapes (Fig. 1). We categorized fibroblasts with PRM1 expression into two nuclear morphologies: “straight” (Fig. 1R) and “curved/hoocked” (Fig 1. S-T). Nuclei classified as “hooked” display two poles that converge and do not align with a tangent passing through the nucleus center. Conversely, “straight” nuclei lack both convergence and divergence of the poles and align along a linear axis intersecting the nucleus center.

**Figure 1.**
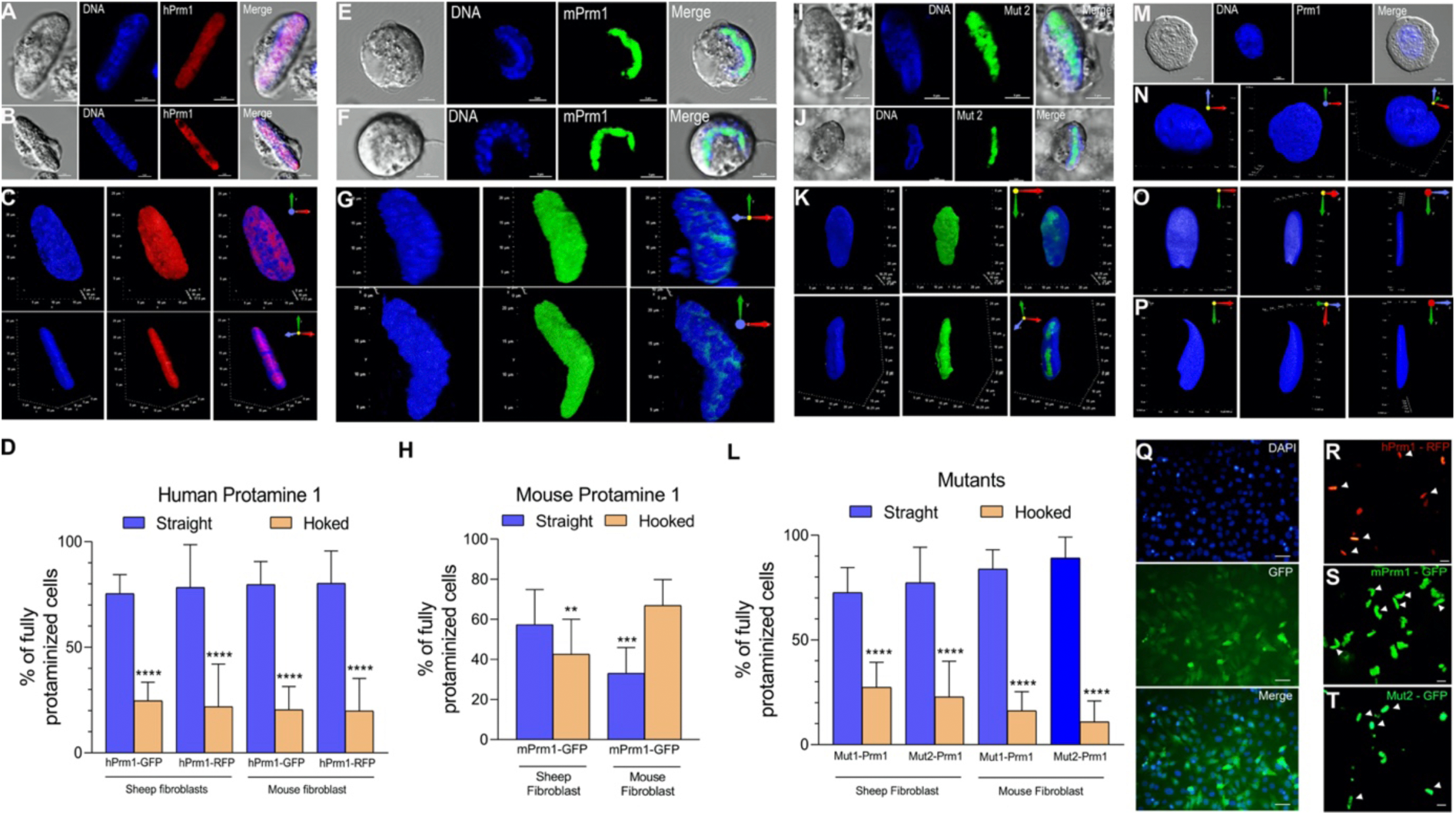
Remodeling of somatic cell nucleus driven by protamine 1 (PRM1). (A) Representative photo of two-dimensional structure of sheep (A) and mouse (B) fibroblasts transfected with human protamine 1 tagged with RFP (*hPrm1*-RFP); Left – bright field, blue – nuclei, cells stained with Hoechst 33342; red: nuclei with *hPrm1*-RFP. Scale bar: 5 µm. (C) Three-dimensional structures of straight somatic nucleus in fibroblasts expressing *hPrm1*-RFP; front view (on the top), side view (on the bottom); (D) Percentages of straight and curved (hooked) nuclei in sheep and mouse fibroblasts expressing human protamine vector (** denotes P<0.05, ****denote P < 0.0001 for differences between straight and hooked nuclei); (E) Representative photo of two-dimensional structure of sheep (E) and mouse (F) fibroblasts transfected with mouse protamine 1 tagged with GFP (m*Prm1*-GFP). Left column – bright field, blue – nuclei, cells stained with Hoechst 33342; green: nuclei with m*Prm1*-GFP. Scale bar: 5 µm; (G) Three-dimensional structures of hooked somatic nucleus in fibroblasts expressing m*Prm1*-GFP; front view (on the top), side view (on the bottom); (H) Percentages of straight and curved (hooked) nuclei in sheep and mouse fibroblasts expressing mouse protamine vector (** denotes P<0.05, ****denote P < 0.0001 for differences between straight and hooked nuclei); Representative photo of two-dimensional structure of sheep (I) and mouse (J) fibroblasts transfected with Mut2-Prm1 tagged with GFP (Mut-2-*Prm1*); Left column – bright field, blue – nuclei, cells stained with Hoechst 33342; green: M*ut2-Prm1-GFP*. Scale bar: 5 µm. (K) Three-dimensional structures of straight somatic nucleus in fibroblasts expressing Mut2-*Prm1*; front view (on the top), side view (on the bottom); (L) Percentages of straight and curved (hooked) nuclei in sheep and mouse fibroblasts expressing mutated protamine vector (** denotes P<0.05, ****denote P < 0.0001 for differences between straight and hooked nuclei); (M) Two-dimensional structure of somatic cells (control - non protaminized); (N-P) Three-dimensional structures of somatic (N) sheep (O) mouse (P) and sperm nuclei in front view (left), side view (right) and rotated 45° (centre); (Q) Negative control group: fibroblasts transfected with a GFP-expressing plasmid lacking the Prm1 gene. Scale bar: 20 µM, (R-T) Representative image illustrating the appearance of ‘fully protaminized’ cells (arrowheads) exhibiting either the straight (R) or hooked (S, T) nuclear morphology, following transfection with hPrm1-GFP, Mut2-GFP, and mPrm1-GFP, respectively. Scale bar: 10 µm.

These categories reflect the distinct nuclear condensation states observed at the fully protaminized stage (Fig. 1R-T). Only cells that exhibited clear nuclear condensation - indicating successful protamine incorporation - were included in this analysis. Result shown, that when human PRM1 (hPRM1) was expressed in sheep (Fig. 1A) or mouse (Fig. 1B) fibroblasts, the nuclei largely adopted a compact, straight shape (Fig. 1A-C). As shown on Fig. 1D, over 80% of those cells exhibited a straight nuclear morphology (Fig. 1A-D). In contrast, expression of mouse PRM1 (mPrm1) predominantly led to curved/”hooked” nuclei both in sheep (Fig. 1E) or mouse (Fig. 1F) fibroblasts. This effect was particularly observed in mouse fibroblasts, where over 70% of cells displayed hooked shape (Fig. 1H). The remodelled nuclear shapes closely resembled the mature sperm nuclear morphology typical of each species (Fig. 1M-P and Fig. 2). These observations were closely tied to PRM1 expression, as expression of the tag proteins (GFP or RFP) alone had no impact on nuclear remodeling (Fig. 1Q).

**Figure 2.**
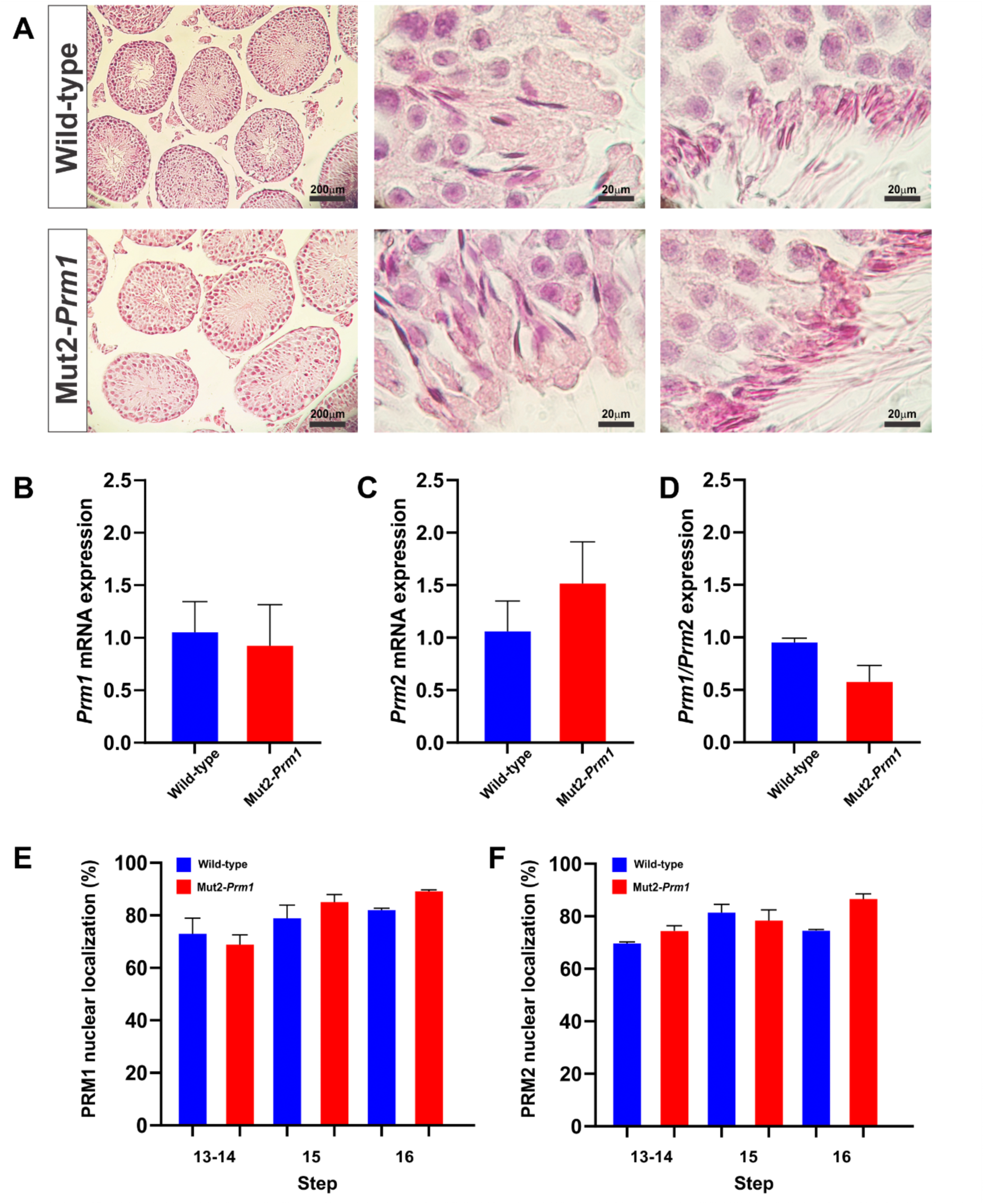
Mutation of PRM1 in cysteine residues (C15 and C29) does not alter spermiogenesis or protamine expression. (A) Testes from adult wild-type (N=3) and Mut2-*Prm1* (N=3) mice were collected for histological studies. Sectioned slides were stained with hematoxylin and eosin and evaluated using a bright-field microscope. No differences were observed between control and Mut2-*Prm1* testes. Representative images for wild-type and Mut2-*Prm1* are shown in the figure. (B) *Prm1* gene expression in wild-type and Mut2-*Prm1* mice (N=3). (C) *Prm2* gene expression in wild-type and Mut2-*Prm1* mice (N=3). (D) *Prm1/ Prm2* mRNA ratio. (E) Percentage of PRM1 localized in the nucleus (N=3) at different spermatid stages during spermiogenesis (from step 13 to 16). (F) Percentage of PRM2 localized in the nucleus (N=3) at different spermatid stages during spermiogenesis (from step 13 to 16). Data are presented as means ± SEM and were analyzed using t-tests for two groups comparison and One-way ANOVA for multiple comparison with GraphPad Prism 10.2.2 software (GraphPad, San Diego, CA, USA).

This observation led us to hypothesize that protamine 1 (PRM1) may underlie these morphological differences. To investigate this, we compared the amino acid sequences of mouse and human PRM1, noting a relatively low sequence homology of 62% (Supplementary Fig. 1A). One notable difference is the presence of three additional cysteine residues in mouse PRM1. Given that cysteines can form disulfide bonds, stabilizing protein structures, these residues may influence chromatin compaction and potentially contribute to the distinct hooked nuclear shape observed in murine sperm.

To test the role of these cysteines, we performed site-directed mutagenesis to replace each additional cysteine. In the first mutant vector (Mut1-Prm1), we replaced cysteine at position 15 with tyrosine (C15Y), which aligns with the human sequence. We also substituted glycine at position 44 with alanine and cysteine at position 45 with tyrosine. In the second mutant vector (Mut2-Prm1), we introduced an additional substitution by replacing cysteine at position 29 with serine, resulting in a double-mutant form (Supplementary Table 1, Supplementary Fig. 1B).

To evaluate the effects of these cysteine modifications on nuclear shape, we transfected both sheep and mouse fibroblasts with either Mut1-Prm1 or Mut2-Prm1. Transfection of sheep fibroblasts with Mut1-Prm1 resulted in a straight nuclear morphology in approximately 70% of cells (Fig. 1L), while Mut2-Prm1 increased this proportion to 77.3% (Fig. 1I-L). These findings support the hypothesis that cysteine residues in mouse PRM1 contribute to the species-specific curved nuclear morphology, and that modifying these residues induces a straighter nuclear shape, more akin to human sperm.

### Mutation of PRM1 cysteine residues (C15, C29) does not affect male fertility

In order to test our hypothesis of the critical role of C15 and C29 in imposing the mature sperm nuclear shape, we genetically modified mice using CRISPR/Cas9 system to explore the *in vivo* function of cysteine residues (C15 and C29) of mouse protamine 1 in sperm DNA condensation and packaging, fertility and sperm head morphology. A Mut2-*Prm1* mutant was generated by intercrossing the founder individuals and the progeny of mice used for further breeding for at least three generations. Mut2-*Prm1 m*ale mice were fertile and sired offspring with an average litter size of 5.8 pups/litter, which is within the range of the reproductive performance of wild-type C57BL/6 mice 7.2 pups/litter (Verley et al., 1967; Nagasawa et al., 1973). There were not observed statistical differences between WT and Mut2-Prm1 mice, with averages of 7.2 pups/litter and 5.8 pups/litter, respectively (P = 0.0797 - based on 17 litters). This indicates that the mutations in cysteine residues (C15 and C29) of PRM1 did not affect reproductive performance in these mice.

Histological analysis of the testes showed no alterations in the seminiferous tubules, which contained the expected balance of germ cells (Fig. 2A). Additionally, there were no distinguishable differences between the spermatids from Mut2-*Prm1* and wild-type adult mice (n=3 each), suggesting no alterations during the spermiogenesis process (Fig. 2A). Next, levels of protamine expression were investigated in wild-type and mutant mice. RNA samples used for gene expression had optimal purity (A260/280>2). There were no significant differences in the expression of protamine mRNA in wild-type (*Prm1*: 1.07 ± 0.29; *Prm2*: 1.07 ± 0.29) and Mut2-*Prm1* mice (*Prm1*: 0.94 ± 0.39; *Prm2*: 1.53 ± 0.39) (P>0.05) (Fig. 2B-C). The ratio between *Prm1* and *Prm2* was also not significantly different between groups (WT: 0.97 ± 0.04 vs. Mut2-*Prm1*: 0.59 ± 0.16; P>0.05; Fig. 2D). These results suggest that *Prm1* mutation causes no major changes in the expression of *Prm1* or *Prm2* mRNA.

In the final stages of spermiogenesis, protamines are transported from the cytoplasm to the nucleus, where they replace histones to facilitate chromatin condensation (Balhorn, 2007; Meistrich et al., 2005). This process is critical for sperm development and function. To investigate whether mutations in the cysteine residues (C15 and C29) of PRM1 impact the nuclear transport of this protein, subcellular immunolocalization of protamines was examined (Supplementary Figure 2). Quantitative analysis showed no significant differences between wild-type and Mut2-*Prm1* mice in the percentage of either PRM1 or PRM2 localized within the nucleus (Fig. 2E-F). These findings suggest that the mutations in the cysteine residues of PRM1 do not interfere with its transport to the nucleus during spermiogenesis.

### Substituting cysteine residues in the PRM1 sequence results in modifications in sperm head morphometry

To further investigate the phenotype of Mut2-*Prm1* mice, we examined the morphological characteristics of their spermatozoa. Spermatozoa from WT and Mut-*Prm1* mice were examined using microscopy and the proportion of normal sperm and with abnormalities in each component (head, midpiece, principal plus terminal piece) quantified (Supplementary Figure 3. We found that, when compared to WT males, Mut2-*Prm1* mice only exhibited a slight increase in the percentage of head abnormalities (P=0.0586; t=2,623, df=4) (Table 1). The proportion of abnormalities in the flagellum (midpiece and principal piece) did not significantly differ between WT and Mut2-*Prm1* mice. Proportions of abnormal sperm were in the range reported previously for adult C57BL/6 mice (Krzanowska 1981, Bruner-Tran et al. 2014, Sekine et al. 2021).

**Table 1.** Percentage of sperm abnormalities in wild type and Mut2-*Prm1* mice. Results are means ± SD. A total of 100 spermatozoa per male (N=3 wild type; N=3 Mut2-*Prm1*) were assessed.

|  | Normal (%) | Head (%) | Midpiece (%) | Principal piece (%) | Other (%) |
| --- | --- | --- | --- | --- | --- |
| Wild-type | 45.7 ± | 2.7 ± 1.52 | 46.7 ± 8.32 | 3 ± 2.0 | 2 ± 3.46 |
| Mut2-*Prm1* | 44.3 ± | 11.0 ± 6.0 | 37.66 ± 10.69 | 5 ± 3.6 | 2 ± 2.64 |

Nuclear morphology was analyzed using methods and software developed by Skinner et al. (2019). This software automatically identifies nuclei and locates key landmarks for orientation and measurement. The nuclear shape is then described using internal angles around the perimeter, generating an “angle profile” (see Methods section for details of methodology).

Overall, results showed minor shape differences between sperm from wild type and from Mut2-*Prm1* homozygous mice using this tool (Figs. 3A-C). Highest variation in the angle profile was seen around position 350 of the profile (Fig. 3C) which corresponds to the flagellum attachment region and the nuclear basal region, as validated previously (Skinner et al. 2019, Skinner 2020). The angle profiles identified five segments in the perimeter of sperm nuclei from of wild-type and Mut2-*Prm1* mice (Fig. 3D). The lentgh of the three of these segments exhibited statistically significant differences, with lower values in Mut2-*Prm1* nuclei in comparison to those of their wild-type counterparts (Table 2).

**Figure 3.**
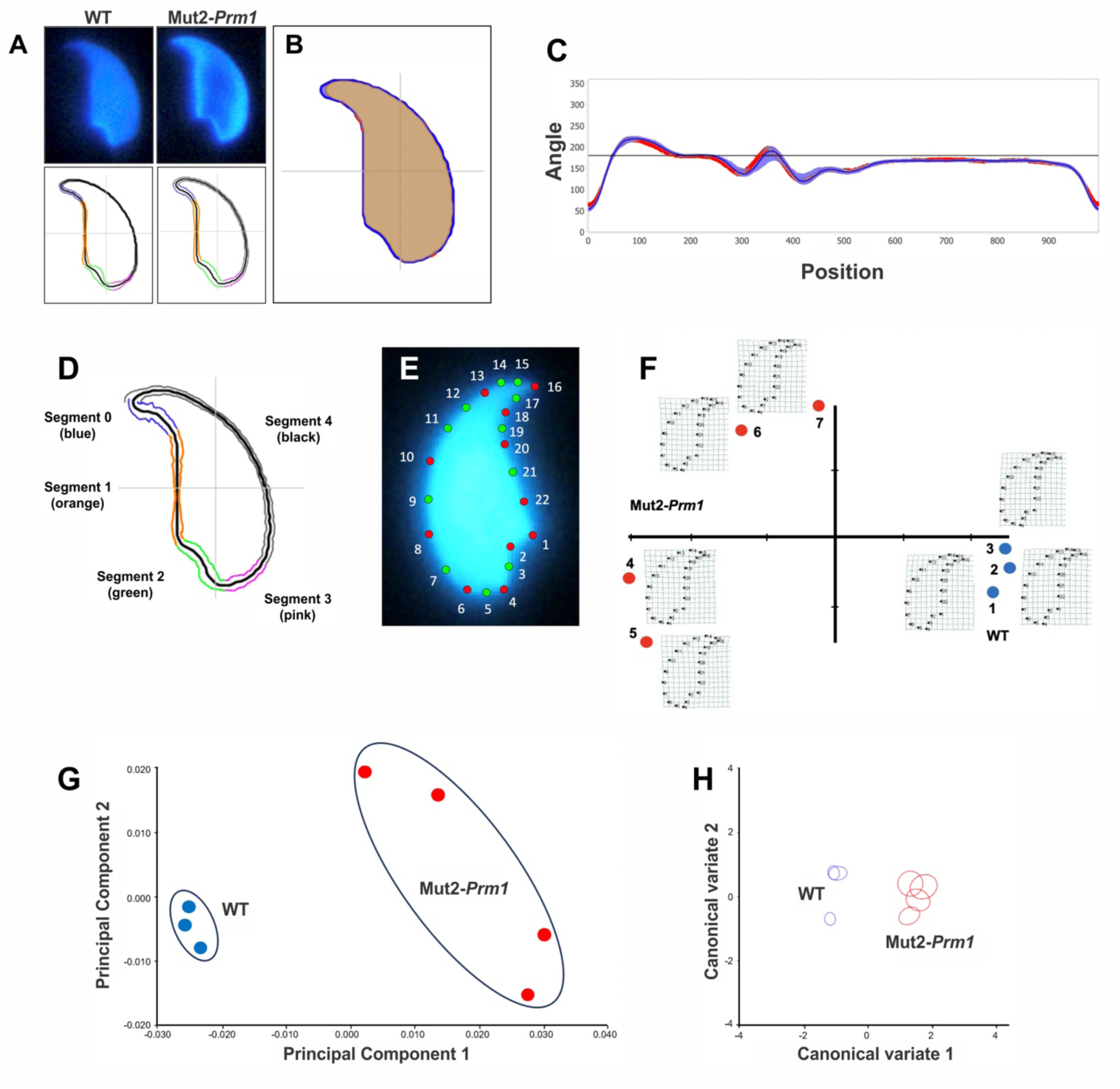
**Changes in PRM1 sequence leads to modifications in sperm nuclear morphology,** as revealed by nuclear morphology analysis (A-D) and geometric morphometrics analysis (E-H). Nuclear morphology analysis (NAM) was carried out using NAM software (Skinner et al, 2019; Skinner 2020) on wild-type sperm (WT; n=3 males; 228 spermatozoa) and Mut2-*Prm1* sperm (n=4 males; 238 spermatozoa). (A) Sperm nuclei from WT and Mut2-*Prm1* mice stained with Hoechst 33258 (above) and consensus nuclei showing segments in nuclear perimeter (below). (B) Superimposition of consensus nuclei from WT (in blue) and Mut2-*Prm1*(in red) sperm. (C) Comparison of angle profiles between WT (blue) and Mut2-*Prm1* (red) sperm showing the median and interquartile range of the nuclear angle profiles. Highest variation is seen around position 350 which corresponds to the flagellum attachment region and the nuclear basal region, as validated previously (Skinner et al, 2019; Skinner 2020). (D) Consensus nuclei from wild type sperm showing segments. Dimensions of each segment for WT and Mut2-*Prm1* sperm are given in Table 2. (E) Geometric morphometrics analysis was carried out as described in the Methods section on sperm from wild type (N=3 males; 261 spermatozoa) and Mut2-*Prm1*) (N=4 males; 196 spermatozoa) animals. Mouse sperm nucleus stained with Hoechst 33342 exhibiting position of landmarks (red) and semilandarks (green) used for geometric morphometric analyses. (F) “Relative warps” analysis with consensus shapes in deformation matrix; WT males are shown using blue symbols, Mut2-*Prm1* males are shown using red symbols). (G) Principal component analysis with consensus shapes for each group. Each dot corresponds to an individual male in each group; blue, WT; red Mut2-*Prm*. (H) Canonical variate analysis for each male; blue, WT; red Mut2-*Prm*.

**Table 2.** Angle profile segment measurements (in microns; means ± SEM) in wild type sperm (WT) (N=3 males, 228 spermatozoa) and Mut2-*Prm1* sperm (N=4 males; 238 spermatozoa). Asterisks indicate differences between groups (P<0.01).

| Segment | WT | *Mut2-Prm1* |
| --- | --- | --- |
| 0 | 2.00 ± 0.03 | 1.89 ± 0.04* |
| 1 | 4.76 ± 0.10 | 4.66 ± 0.11* |
| 2 | 2.73 ± 0.10 | 2.73 ± 0.09 |
| 3 | 2.01 ± 0.09 | 1.84 ± 0.08* |
| 4 | 10.76 ± 0.17 <sup>a</sup> | 10.80 ± 0.20 |

The nuclear morphology software also provides measurements of various sperm nuclear parameters (Supplementary Figure 4, Supplementary Table 2). Significant differences in sperm head dimensions were noted between wild type and Mut2-*Prm1* mice (Table 3). Differences were observed between wild-type and Mut2-*Prm1* sperm in all linear dimensions with the exception of bounding height and width of body. Linear dimensions were lower in Mut2-*Prm1* in comparison to wild type sperm. Dimension-derived parameters that infer shape followed two patterns: ellipticity and elongation were higher in Mut2-*Prm1* sperm whereas aspect ratio and regularity were lower, in comparison with wild-type sperm. Circularity was not very different between the two types of sperm. These results indicate that sperm nuclei from Mut2-*Prm1* mice were more elongated (streamlined) and less regular in shape than those from wild-type mice.

**Table 3.** Morphometry of sperm nuclei of wild-type sperm (N=3 males; 228 spermatozoa) and Mut2-*Prm1* sperm (N=4 males; 238 spermatozoa). Measures (in microns) are means ± SEM. Asterisks indicate differences between WT and Mut2-*Prm1* (P<0.01 for all differences with the exception of minimum diameter for which P<0.05).

| Parameter | WT | *Mut2-Prm1* |
| --- | --- | --- |
| Number of nuclei analyzed | 228 | 238 |
| Area | 21.26 ± 0.12 | 20.52 ± 0.18* |
| Perimeter | 21.89 ± 0.11 | 21.55 ± 0.17* |
| Maximum feret | 8.21 ± 0.03 | 8.07 ± 0.04* |
| Minimum diameter | 3.43 ± 0.02 | 3.32 ± 0.03* |
| Bounding height | 7.34 ± 0.03 | 7.34 ± 0.05 |
| Bounding width | 5.48 ± 0.04 | 5.15 ± 0.05* |
| Width of body | 3.78 ± 0.04 | 3.70 ± 0.03 |
| Length of hook | 1.74 ± 0.03 | 1.49 ± 0.04* |
| Circularity | 0.56 ± 0.00 | 0.57 ± 0.01* |
| Ellipticity | 1.36 ± 0.01 | 1.46 ± 0.02* |
| Aspect ratio | 0.75 ± 0.01 | 0.71 ± 0.01* |
| Elongation | 0.15 ± 0.00 | 0.18 ± 0.01* |
| Regularity | 1.48 ± 0.01 | 1.44 ± 0.01* |

Using geometric morphometrics analysis, Procrustes ANOVA revealed significant differences (P<0.0001) in both size (centroid size) and form between wild-type and homozygous Mut2-*Prm1* spermatozoa (Fig. 3E-F). This indicates notable variations in shape between the two groups, suggesting an impact of the mutation on sperm head conformation. Principal Component Analysis clearly distinguished between the two groups (Fig. 3G). To assess the significance of the separation between subpopulations based on Procrustes distances, canonical variable analysis was conducted. Significant distances (P<0.05) were noted between any group of mutated individuals and the wild-type groups. Ellipses representing 90% confidence with respect to the consensus are depicted (Fig. 3H).

### Cys15 and Cys29 mutations in mouse protamine 1 affects chromatin condensation and histone retention

Transmission electron microscopy (TEM) unveiled that the mutation of cysteines 15 and 29 in the protamine sequence disrupted chromatin condensation, resulting in a looser structure with multiple low-density foci scattered throughout the nuclei. This phenotype, absent in mouse control spermatozoa, resembles the ultrastructure of human sperm nuclei, characterized by nuclear domains retaining a histone organization (Fig. 4A). To delve deeper into this observation, Western blot analysis was performed on sperm from wild-type and Mut2-*Prm1* mice. Remarkably, a high level of histone H3, as an indicator for histone retention, was detected in Mut2-*Prm1* sperm (Fig. 4B), consistent with the presence of low-density foci. In wild-type mice, only a minimal fraction (1%) of nucleosome-organized domains is typically observed (Brunner et al., 2014), while in human spermatozoa, histone retention accounts for up to 10-15% of the sperm DNA (Hammoud et al., 2009; Yamaguchi et al., 2018). These findings underscore the influence of PRM1 cysteines on overall histone retention, potentially contributing to reduced DNA condensation.

**Figure 4.**
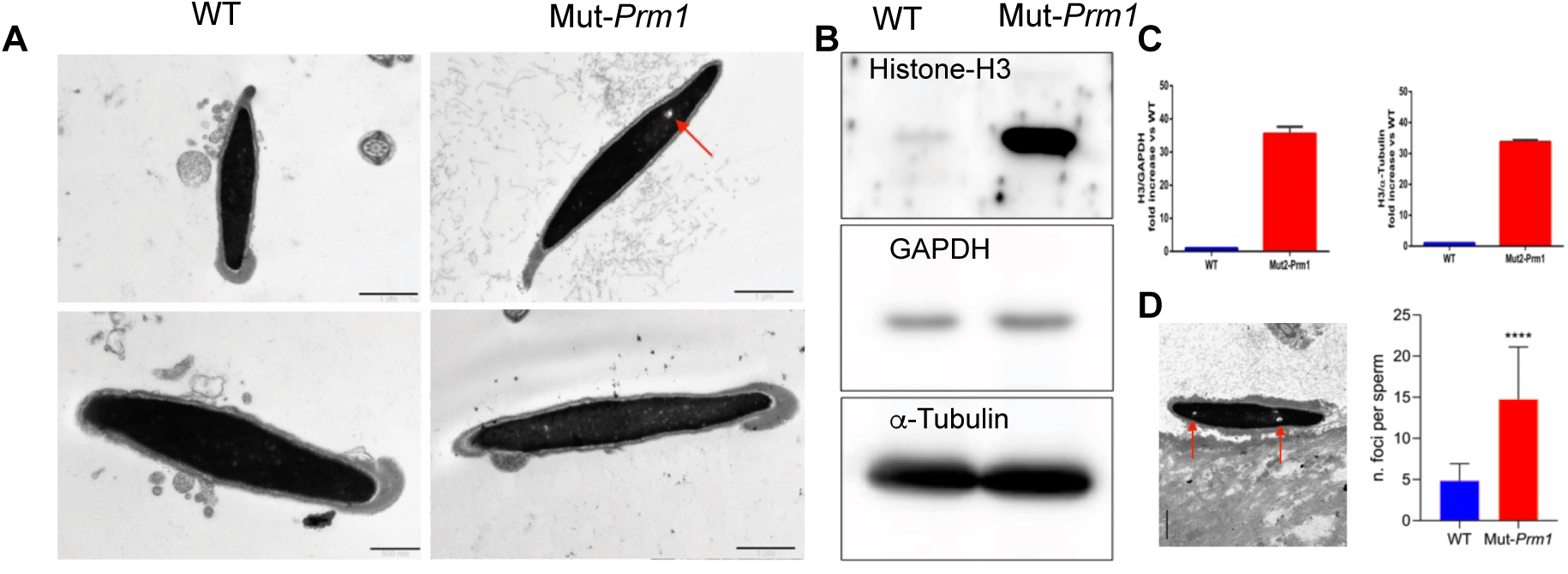
**Electron microscopy and histone retention** of wild-type (WT) and Mut2-*Prm1* sperm. (A) Transmission electron micrographs (TEM) of wild-type (WT) and Mut2-*Prm1* sperm. Note reduced chromatin condensation (red arrows). Scale bar = 1 µm or 500 nm (upper right panel). (B) Immunoblot against histone H3, GAPDH and tubulin of basic nuclear proteins of sperm from wild type (WT) (N=3) and Mut2-*Prm1* mice (N=3). (C) Quantification of Western blot bands by densitometry; top: density of histone-H3 in relation to GAPDH; bottom: density of histone-H3 in relation to *a*-tubulin. (D) Representative transmission electron microscopy (TEM) image of Mut2-Prm1 sperm. Scale bar = 1 µm. The number of foci per sperm is quantified and presented in the accompanying graph. Statistical analysis indicates a significant difference (p < 0.0001).

## Discussion

In this study, we have shown that protamine sequence affects sperm head shape in a simplified *in vitro* model of nuclear remodeling in fibroblasts, and *in vivo* in mutated mouse spermatozoa. This study was motivated by the observation of different nuclear shapes following the forced expression of different PRM1 species (mouse and human). The idea emerged that the difference in the number and position of cysteines, specifically the mouse specific C15 and C29, between mouse and human PRM1 sequences could explain the observed phenomena. Our *in vitro* studies unequivocally demonstrated a link between protamine sequence and nuclear morphology after compaction, a trait that we have further demonstrated by site-directed mutagenesis of cysteines in mouse PRM1 at position 15 and 29, that brings the mouse sequence closer to the human one. The nuclear remodeling by the mutant PRM1 confirmed our prediction that the lack of cysteine 15 would abolish protamine looping resulting from the first intra-molecular 7-15 S-S bond (Suppl. Fig. 1B), which in turn would influence nuclear shape by modifying the way it would associate with DNA and interact with its neighbors, as proposed in the electrostatic zipper model, with a looser condensation capacity (Kornyshev et al. 1998, Sitk et al. 2003). The transfection experiments of fibroblasts with Mut2-*Prm1* also led to an increased number of nuclei with a straight shape similar to the one obtained with the *hPrm1* vector (Fig. 1A, B, E, J).

To verify our conclusion in relevant physiological models, we used gene editing to replace C15 and C19 in the mouse PRM1-encoding gene. Careful nuclear morphology examination with an ad-hoc software (Skinner et al., 2019) revealed differences in the majority of sperm measurements between wild type and Mut2-*Prm1* sperm as well as in dimension-derived parameters inferring shape (Table 3). Further analysis using geometric morphometrics, as developed for mouse sperm (Varea et al. 2013), revealed clear distinctions between wild-type and mutant mice, reinforcing the idea that changes in PRM1 sequence impacts on nuclear sperm shape. The ultrastructural analysis showed differences in chromatin condensation and confirmed morphological changes in mutated spermatozoa. Changes affected the general outline of the cells and thickness of the nuclei (Fig.). It is plausible that in a flattered head, such as those of mouse, sperm with a reduced nuclear compaction would be affected mildly, albeit significantly, in the profile and thickness of the nucleus in mutated sperm. The changes we observed in shape and morphometry of PRM1 mutated sperm in comparison to wild-type controls are in agreement with the observation that sperm nuclear morphology is altered in mice in which the *Prm1* is deficient or is absent (Arévalo et al., 2022; Merges et al. 2022). Interestingly, a mouse with a lysine-to-alanine mutation (K49A) in PRM1 exhibited sperm structural changes, including abnormal head morphology, in the absence of effects on weight, germ cell populations, sperm counts or PRM1 levels (Moritz et al. 2023). These results emphasize the likely role of specific PRM1 residues in nuclear shaping.

It was somewhat unexpected to find altered histone retention in spermatozoa with mutated PRM1. Western blot analysis revealed a marked increase in histone retention (Figure 4B, C). This increased histone retention may contribute to the observed looser nuclear compaction in spermatozoa from the mutated mice, as indicated by the presence of low-density areas in TEM images (Figures 4A). These low-density foci are consistent with previous studies suggesting that disruptions in chromatin compaction, such as those caused by altered protamine-histone interactions, can result in regions of reduced density within the sperm nucleus (Balhorn, 2007; Ward & Coffey, 1991; Oliva & Dixon, 1991). This loosening of chromatin likely underlies the morphometric changes observed in the sperm heads (Figure 3), further supporting the link between histone retention and chromatin decondensation.

Our results provide strong evidence to suggest that, in addition to actions exerted by extranuclear structures, such as the manchette, in shaping the sperm nucleus (Kierszenbaum et al. 2004, Yoshida et al. 2018, Hermo et al. 2010, Russell et al. 1991, Lehti et al. 2016) intranuclear factors are also important for changes in nuclear morphology. In our *in vitro* studies, nuclear remodeling occurs in the absence of a manchette, indicating that the inter/ intra-protamine S-S bonds are sufficient to remodel the nucleus, and that differences in protamine sequences may result in varying nuclear shapes in this somatic cell *in vitro* model. This hypothesis may also be supported by the relatively low conservation of the protamine gene across mammals, a surprising finding for a gene with such a highly conserved function. Apparently, evolution of transition proteins and protamine sequence and their regulation seems to have exerted a major influence in sperm head shape and size (Lüke et al. 2014, Lüke et al. 2016).

In conclusion, we have demonstrated that protamine sequence is an important factor in intrinsic nuclear remodeling and, in particular, that protamine cysteines 15 and 29 play a major role in configuring nuclear morphology, histone retention and chromatin condensation. Because nuclear compaction/remodeling is crucial for male fertility, our results help to characterize mechanisms underlying sperm nuclear formation and to identify the evolutionary forces that influence changes in sperm shape.

## Methods

### Ethics statement

Animal experimentation was regulated and approved by the Italian Ministry of Health, upon the presentation of the research description prepared by the ethics committee of the “Istituto Zooprofilattico Sperimentale di Teramo” (Prot. 944F0.1 del 04/11/2016). The authorization number granted by the Italian Ministry of Health is n° 200/2017-PR. All methods were performed in accordance with the relevant guidelines and regulations.

### Reagents

Unless otherwise stated, all materials used were purchased from Sigma.

### Plasmid construction

The plasmids containing the GFP-tagged sequence of mouse *protamine 1* gene (m*Prm1*) and GFP or RFP-tagged sequence of human *protamine 1* gene (h*Prm1*) were obtained as described previously (Czernik et al. 2016).

### Site-Directed Mutagenesis of Mouse Prm1: “Humanization” of the Mouse Protamine Plasmid

Human and mouse protamines share a relatively low homology (62%, Supplementary Fig. 1A), with mouse PRM1 containing three additional cysteine residues. To investigate the impact of protamine sequence on nuclear remodeling, we used site-directed mutagenesis to alter these cysteine residues.

In the first mutated vector (Mut1-Prm1), we replaced Cys15 with Tyr, corresponding to the amino acid found in human PRM1 (C15Y; Supplementary Table 1, Supplementary Fig. 1A). Specifically, nucleotide 44 was altered from Gly to Ala, and nucleotide 45 was modified from Cys to Tyr, introducing a change that reduces the cysteine count to two more than the human counterpart.

For the second mutated vector (Mut2-Prm1), we incorporated an additional mutation. Besides the above modifications, nucleotide 85 was changed from Tyr to Ala, resulting in an exchange of Cys for Ser at codon 29. With this mutation, the plasmid’s protamine has only one cysteine more than human PRM1 (Supplementary Table 1, Supplementary Fig. 1A).

These mutations were achieved via in vitro site-directed mutagenesis using the QuikChange Multi-Site Directed Mutagenesis Kit (Agilent Technologies), with primers designed using Agilent’s tool (Supplementary Table 1). Sequence analysis confirmed the successful generation of both Mut1 and Mut2 plasmids.

### Fibroblast transfection

Sheep fibroblasts (SFs) and mouse fibroblasts (MFs) were cultured in a petri dish in Dulbecco’s Modified Eagle Medium (DMEM) (GIBCO) containing 2 mM glutamine, 3.7 g/L NaHCO_3_, 0.5% (v:v) gentamicin and 10% (v:v) Fetal Bovine Serum (FBS) at 38.5°C (SFs) or 37°C (MFs) under 5% CO_2_/air in a humid atmosphere. At passage 4-6 cells were transfected with plasmids m*Prm1*-GFP, h*Prm1*-RFP, pGFP-*Prm1* g44a c45t and pGFP-*Prm1* g44a c45t, t85a as previously described (Czernik et al. 2016) with some minor modifications (Palazzese et al. 2018). Briefly, SFs and MFs at 80% confluence were starved for 24 h in DMEM medium supplemented with 0.5% (v:v) FBS. At 4 h post-transfection, cells were treated with 50 nM Trichostatin A (TSA) - histone acetylation using histone deacetylase inhibitor (HDAI) - for 16 h, to encourage the chromatin opening and, progressively, the protamine incorporation into the DNA.

### Evaluation of nuclear morphology in transfected fibroblasts

At 48 h post-transfection cells were stained with 5 µg/ml of Hoechst 33342 for 10 min and analysed under confocal microscopy (Nikon Eclipse Ti-E) using NIS-Elements Confocal software (Nikon). To evaluate the nuclear morphology only the cells that have reached the full protaminization stage (i.e., a spermatid-like condensed nucleus) were analysed. The nuclear shape was categorized as “straight” or “hooked” (“curved”). The chromatin conformation rate (straight or hooked) was calculated in relation to the total number of spermatid-like cells.

### Generation of protamine 1 mutant mice

A mouse line with the mutated protamine (Mut2-*Prm1*) was generated by the injection of CRISPR/Cas9 to zygotes, using two sgRNAs in C57BL/6 background. Exclusvely homozygotes male have been used in all analysis. Introduced modifications includes: the insertion of missense c.44G>A, c.45C>T mutation; EcoRV (Eco32I) restriction site for rapid routine genotyping; 3 silent mutations (c.48C>G, c.49C>A, c.51T>A) to prevent the generation of chimeric mice by CRISPR/Cas9 system and unwanted 21 bp deletion in the end of exon1 (c.del85_105delTGCTGCCGGCGGAGGAGGCGA) spanning over 7 amino acids (p.24_30delArgArgArgAgrArgCysCys). Localization of insertion is based on Ensembl Prm1 Prm1-201 transcript ENSMUST00000023144.6

Left Target gRNA sequence [PAM sequence]: GACATCTTCGCCTGCGACGG [CGG]

Right Target gRNA sequence [PAM sequence]: ACTTACTTCGCCTCCTCCGC [CGG]

ssDNA repair template (5’-3’) was synthetized by annealing two long oligonucleotides, single-stranded overhangs were filled with PCR.

Donor dsDNA to introduce the mutation was subsequently reamplified using short primers and reverse/complement strand was removed. Sequence of donor dsDNA: TGCTCTGAGCCAGCTCCCGGCCAAGCCAGCACCATGGCCAGATACCGATGCTGCCGCAGCAAAAGCAGGAGCAGATATCGGAGACGCAGGCGAAGATGTCGCAGACGGAGGAGGCGAAGCTGCCGCCGGAGGAGGCGAAGTAAGTAGAGGGCTGGGCTGGGCTGTGGGGGGTGTGGGGTGCGGGACTGGGCAGTCT.

### Genotyping

Samples for genotyping were collected from one month old PRM1 mutant mice by ear punch. Each sample was divided in two to secure a backup. ***DNA isolation:*** To each sample 75 µl of 25 mM NaOH / 0.2 mM EDTA (Alkaline Lysis Reagent) was added. Next, samples were placed in thermocycler at 95°C for 1 h, then the temperature was reduced to 12°C until ready to proceed to the next step. Then, 75 µl of 40 mM Tris HCl (pH 5.5) was added to the samples followed by centrifugation at 4000 rpm for 3 min. This was used as a DNA aliquot for PCR reaction. ***PCR reaction:*** PCR reaction mix contained (per sample): 12.5 µl Color Taq PCR Master Mix (2×) (EURx Ltd, Gdańsk, Poland; Cat. #E2525); 0.4 µl for each primer; 10.7 µl H_2_O and 1 µl of DNA sample. PCR conditions were an initial denaturation step of 95°C for 3 min, followed by 35 cycles of 95°C for 45 sec, 60°C for 30 sec, 72°C for 1 min and a final extension at 72°C for 7 min. After amplification was completed, 10 µl of PCR product was loaded onto 1.5% agarose gel containing 1X TAE (Tris-acetate-EDTA) buffer (Cat. #15558042; Invitrogen™, Waltham, USA) and stained with 5 µl of SimplySafe™ per 100 ml of agarose solution (Cat. #E4600-01; EURx Ltd, Gdańsk, Poland). Electrophoresis was performed at 100 V for 30-45 min with ladder for sizing and visualized by ChemiDoc XRS+ System (Bio-Rad) and Image Lab Software.

#### Primer sequences

PRM1-F 5’-CAGGCCACAGCCCACAAAAT-3;

PRM1-R 5’-TTGAAGTCTGGTAAAATTCTCACGC-3’ (product size: 403 bp).

#### Restriction enzyme digestion

Reaction was performed using FastDigest Eco32I (Thermo Scientific; Cat. # FD0303; Waltham, USA). Reaction mix contained (per sample): 10 µl DNA, 2 µl buffer, 17 µl H_2_O, 1 µl enzyme. Samples were incubated 30 min at 37°C. After the incubation was completed, 20 µl of sample was loaded onto 1.5% agarose and electrophoresis was conducted and visualized as above. PCR products sizes after digestion were corresponding to: 403 bp – WT; 403 bp + 253 bp + 150 bp – Heterozygote; 253 bp + 150 bp - Homozygote.

### Testis protamine expression

RNA was extracted from testes (N=3 for wild type; N=3 for Mut2-*Prm2* mice) using the E.Z.N.A.® HP Total RNA extraction kit (Omega-Biotek, Norcross, GA, USA) following the manufacturer’s instructions. RNA concentration and purity was assessed with a spectrophotometer (NanoDrop 2000, ThermoFisher, Walthman, MA, USA) and ratios of absorbances A260/280 were obtained. Reverse transcription polymerase chain reaction (RT-PCR) was performed with the Superscript III First-Strand Synthesis kit (Invitrogen, Waltham, MA, USA), following the guidelines and using Oligo-dT primers for the retrotranscription. The iQ™ SYBR® Green Supermix kit (BioRad, Hercules, CA, USA) was used for assessing the relative gene expression of *Prm1* and *Prm2*. Quantitative PCR (qPCR) was carried out in a thermal cycler (QuantStudio™ 3 Real-Time PCR Instrument, 96-well 0.1 ml Block, Applied Biosystems, Waltham, MA, USA). Primers and PCR conditions were as previously described (Lüke et al. 2014). Primers for *Prm1* (NM_013637.5) were: forward [5’-3’], AGGCGAAGATGTCGCAGACG; reverse [5’-3’], CCGCCGCTCATACACCATAAGG. Primers for *Prm2* (NM_008933.2) were: forward [5’-3’], ACAAGAGGCGTCGGTCATGC; reverse (5’-3’), GGTGCAGGAAATGTAGGAGGCAC. Primers for rRNA 18s (housekeeping, NR_003278.3) were: forward (5’-3’), GTGATGGGGATCGGGGATTGCA; reverse [5’-3’], AGAGGGACAAGTGGCGTTCAGC. The cycle for PCR was the following: a 95°C cycle for 10 min, 40 cycles of 15 s at 95°C and a cycle at 62°C for 1 min. A melt-curve was performed at the end of the cycles: 95°C for 15 min, 1 h at 50°C and 15 min at 95°C.

The results of qPCR were evaluated with the 2^-ΔΔC^_T_ method (Livak & Schmittgen, 2008). Briefly, Ct values of the triplicates of each sample and each gene were averaged and the ΔC_T_ (C_T_ PRM – C_T_ 18S) was estimated for each sample. ΔΔC_T_ was calculated by subtracting ΔC_T_ of each sample minus the average of ΔC_T_ 18S. Finally, 2^-ΔΔCT^ was obtained and averaged for each group.

### Testis histological preparation and examination

The testes of adult mice (N=3) were dissected and fixed in a 4% formaldehyde solution in PBS, paraffin-embedded, and sectioned into 5 µm slices. Hematoxylin and eosin staining was carried out using standard procedures. Histology was recorded using a digital camera (Olympus QColor5), attached to an Olympus BH-2 microscope (Olympus, Center Valley, PA).

### Protamine translocation to the nucleus

Staining for the localization of protamines was carried out as described before (Agudo-Rios et al. 2023). Briefly, isolated mixed germ cells from adult wild-type (N=3) and Mut2-*Prm1* (N=3) mice were fixed in 4% paraformaldehyde/PBS (with 0.1 M sucrose) for 15 min at room temperature. After 3 washes with PBS, cells were resuspended in PBS and spread on SuperFrost/Plus slides (Fisher Scientific, Pittsburgh, PA). Slides were blocked with a solution containing 10% goat serum (Vector Laboratories, Inc., Burlingame, CA), 3% BSA (Sigma-Aldrich, St. Louis, MO) and 0.2% Triton X-100 (Fisher Scientific, New Jersey, NY), and incubated at room temperature for 1 h. The primary antibodies (anti-PRM1, Hup1N, or anti-PRM2, Hup2B; Briar Patch Bioscience, Livermore, CA) were diluted 1/100 in the blocking solution and incubated overnight at 4°C. Following incubation, samples were washed 3 times (10 min each) with PBS and incubated with secondary antibody (Alexa Fluor 594-conjugated anti mouse IgG, Jackson ImmunoResearch Laboratories, West Grove, PA) at room temperature (in the dark) for 1–2 h. After 3 washes with PBS, slides were mounted with VectaMount with DAPI mounting media (Vector laboratories, Inc., Burlingame, CA) and sealed with a coverslip and nail polish. Images were captured with a Zeiss LSM700 confocal laser-scanning microscope and analyzed using ImageJ. In order to differentiate spermatids stages, we examined the topological morphology of each cell. This evaluation encompasses the analysis of nuclear positioning within the cytoplasm, the configuration of the acrosome, and the presence or absence of the manchette structure. After the cells were classified in the respective step, the percentage of protamines in the nucleus and the cytoplasm was calculated by measuring the immunofluorescence intensity of PRMs in each compartment. To conduct this, DAPI staining was used to indicate the nucleus and a DIC image was used to determine the total cell area. Fluorescence in the cytoplasmic region was derived by subtracting nuclear fluorescence from the fluorescence in the entire cell, then divided by the respective area to normalize. Subsequently, nuclear values were multiplied by 100 and divided by total cell fluorescence.

### Sperm collection and preparation

Spermatozoa were recovered from females after mating with males (either wild type or Mut-*Prm1*). Female was mated with an appropriate male until a vaginal plug was observed (mating success was checked every hour). Females were sacrificed and the uterus was dissected to recover spermatozoa from one ejaculation. The uterus was placed in a 35 mm Petri dish containing PBS and 0.4% BSA and the contents were flushed using a 1 ml syringe. The sperm suspension was transferred to a 1.5 ml Eppendorf tube and centrifuged at 1500 rpm for 10 min. Supernatant was discarded and fixative was added (2% v/v glutaraldehyde in 0.165M sodium cacodylate/HCl, pH 7.4). Fixation was performed overnight at 4°C. Samples were centrifuged as above, supernatant was discarded and suspended in 0.1 M cacodylate buffer, pH 7.3. Fixed samples were stored at 4°C until analysis.

### Sperm abnormalities

Sperm morphology was evaluated by placing 10 µl of fixed sperm suspension between a slide and a 22×22 mm coverslip (without applying manual pressure). Spermatozoa in the preparation were allowed to settle for 5 min and then examined at 400x and 1000x magnification using phase contrast optics (Kawai et al. 2006). A total of 100 spermatozoa per male (N=3 wild type; N=3 Mut2-*Prm1*) were examined to quantify morphological abnormalities of the head, midpiece and principal plus terminal piece (Kawai et al. 2006, Shin et al. 2009, Oliveira et al. 2010), Bruner-Tran et al. 2014) (Supplementary Figure 3).When a sperm cell exhibited more than one abnormality, the more severe one was recorded to avoid overestimations. The percentage of normal sperm was calculated as the proportion of spermatozoa with no morphological abnormalities out of all spermatozoa examined.

### Nuclear morphology analysis

The Nuclear Morphology Analysis program was employed to measure and compare the shapes and dimensions of sperm nuclei (Skinner et al. 2019, Skinner 2022). The program uses a modified Zahn-Roskies transformation to convert the outlines of objects into linear profiles, with a set of rules to identify landmarks of interest from these profiles, detecting subtle variation in nuclear shape (Skinner et al. 2019, Skinner 2022). The method for morphological analysis allows to automatically identify nuclei in microscopy images, and then find key landmarks for orientation and measurement. An angle profile is generated that describes the shape of the nucleus by measuring internal angles around its perimeter .

Different sperm features (the “hook”, site of flagellum attachment) can be identified in the profile regardless of the orientation of the sperm nucleus in the image. Profiles for different data sets (WT vs Mut2-*Prm1* sperm) were aligned against each other. The software calcules median and interquartile range of the angles which are used to illustrate differences between sets of data. Detailed information about the basis for shape analyses, as well as validation of correspondence between segments and sperm features, are available (Skinner et al. 2019; Skinner 2020; (https://bitbucket.org/bmskinner/nuclear_morphology/wiki/Home). In addition to the shape profiles, several measurements of nuclear parameters are automatically calculated and displayed (Supplementary Figure 4, Supplementary Table 2).

### Nuclear morphology analysis using geometrics morphometrics

Geometric morphometrics methods were used to analyze head shape variation based on a set of landmarks that correspond to the spatial position of particular anatomical traits (Kendall, 1986, Goodall 1991). Analyses did not include sperm with a nuclei that departed from a normal form. A total of 22 bidimensional landmark coordinates were obtained from spermatozoa of wild-type and Mut2-*Prm1* mice. Landmark data were processed as described previously (Varea Sanchez et al. 2013, Luke et al. 2014). All morphometric analyses were conducted with MORPHOJ (Klingenberg 2011).

### Transmission electron microscopy (TEM) and measurements of sperm head thickness

Sperm were collected, as described above, and fixed in glutaraldehyde (2.5% in 0.1 M cacodylate buffer, pH 7.4) for 24 h. After washing in ddH_2_O, cells were post-fixed in 2% OsO_4_ in ddH_2_O for 2 h. Next, cells were dehydrated through a graded series of ethanol solutions (30%, 10 min; 50%, 15 min; 70%, 24 hr; 80%, 10 min; 96%, 10 min; 100%, 10 min; acetone, twice for 15 min) and were infiltrated with graded concentrations of Epoxy 812 Resin (EPON) resin in 100% acetone (1:3, 20 min; 1:1, 24 hr; 3:1, 2 hr), infused twice for 1 hr in pure EPON resin, and polymerized at 65° C for 24 hr. Next, 60-nm sections were prepared and examined using a LEO 912AB electron microscope. Images were captured using a Slow Scan CCD camera (Proscane) and EsiVision Pro 3.2 software (Soft Imaging Systems GmbH).

### Western blot analysis

Sperm collected from Mut2-*Prm1* (N=3) and wild type mice (N=3) were lysed using 8 M urea buffer at 4°C overnight, then samples were sonicated for 20 s each and centrifuged at 13,000 xg for 5 min at 4°C. Protein concentration was assessed using the BCA Protein Assay Kit (Thermo Fisher, Milan, Italy) according to the manufacture’s protocol. For each sample, 50 µg of protein were heated at 95°C for 5 min and then loaded into a 10% SDS polyacrylamide gel. Proteins were transferred onto a 0.45 µm nitrocellulose membrane (Bio-Rad, Milan, Italy) at 4°C for 2 h and 200 mA. After transfer, the nitrocellulose membrane was blocked with 5% non-fat dry milk in 0.1% Tween-20 PBS (PBST) for 1 h at room temperature. Membranes were incubated overnight with primary antibodies rabbit anti-Histone H3 (PTM-1001RM, PTM BIO, Chicago, Illinois, USA), mouse anti-α-Tubulin (DM1A; 3873S, Cell Signaling Technology, Inc., USA) and mouse anti-GAPDH (D4; MA1-16757, ThermoFisher Scientific, USA) diluted 1:1000 in PBST with 0,5% non-fat dry milk. Then, membranes were washed three times for 15 min with PBST and incubated with secondary antibodies HRP-conjugated Affinipure goat anti-mouse IgG(H+L) (SA00001-1, Proteintech, Manchester, UK) and mouse anti-rabbit IgG-HRP (sc-2357, Santa Cruz Biotechnology, USA) diluted at 1:10000 in PBST for 1 h at room temperature. Final detection was performed using enhanced chemiluminescence (ECL) Western Blotting Substrate (Amersham, Pharmacia, Piscataway, NJ, USA) and image acquisition was carried out using the ChemiDoc System (Bio-Rad, Milan, Italy). Western blot analyses were repeated 4 times.

### Statistical analysis

Data were analyzed using GraphPad Prism for Windows (Version 6.01, GraphPad Software, Inc., CA, USA). Statistical analyses of transfection yield and the rates of different nuclear morphologies were based on three to five replicates per experiment and compared using Fisher’s exact test or ANOVAs with transformed data (arcsine for percentages and log10 for other parameters). Statistical analyses of protamine expression were performed by a two-tailed T-test. Statistical analyses of nuclear translocation of protamines was assessed using One-way ANOVA with GraphPad Prism version 10.2.2. The significance level for all statistical tests was set at *P* < 0.05.

## Author contributions

M.C. and L.P. conducted experiments, figure preparation, analyzed data, manuscript writing. D.W animal breeding and sperm collection. D.I. synthesis of mutated expression vectors. S.K. conceived the experiments, advice in vector design, manuscript writing. J.F. and H.F. analyzed data, manuscript preparation. R.V., J.A.R., A.-R., N.L. and C.A.-R. conducted experiments and analyzed data. E.R.S.R. design of the experiments, analyzed data, manuscript writing. M.E.T. conducted experiments, analyzed data, figure preparation, manuscript editing. P.L. design of the experiments, analyzed data, manuscript writing.

## Acknowledgements

The authors warmly acknowledge Dr. L. Valbonetti, University of Teramo, for help in acquisitions of the confocal microscopy images as well as Aurora Scutieri, University of Teramo for Western blot analysis.

## Data and materials availability

All data are available in the manuscript or the supplementary materials.

