## Supplementart for "Protamine sequence determines species-specific nuclear shape and histone retention": Supplementary_Czernik_Nucelar_Shape.pdf

### SUPPLEMENTARY MATERIAL

#### Supplementary Material and Methods

#### Bioinformatics tools

Human and mouse protamine 1 sequences were retrieved from the UniProt database (www.uniprot.org). The sequence alignment between human and mouse protamine 1 was performed with the ALIGN tool in the UniProt portal (www.uniprot.org). Multiple sequence alignment was performed with jalview (Waterhouse et al. 2009). Secondary structure prediction was performed with the PSIPRED web server (Buchan et al. 2013) (http://bioinf.cs.ucl.ac.uk/psipred_new/). To identify the relative position of the substituted residues, a 3D model for the structure of human protamine 1 was retrieved. The available template for human protamine 1 is 1SKN.pdb (Rupert et al. 1998): the DNA binding region of the developmental transcription factor protein skinhead 1 (skn-1) from *C. elegans*. The sequence homology is 46% between the residues 490-526 of the template and 8-51 of the target sequence. No template could be found for mouse protamine 1 structure nor for the two mutants. The 3D model of human protamine with the two mutated residues was retrieved from the Modbase server (https://modbase.compbio.ucsf.edu/) (Pieper et al. 2006). Molecular graphics and analyses were performed with the UCSF Chimera package (Pettersen et al. 2004). Chimera was developed by the Resource for Biocomputing, Visualization, and Informatics at the University of California, San Francisco (supported by NIGMS P41-GM103311).

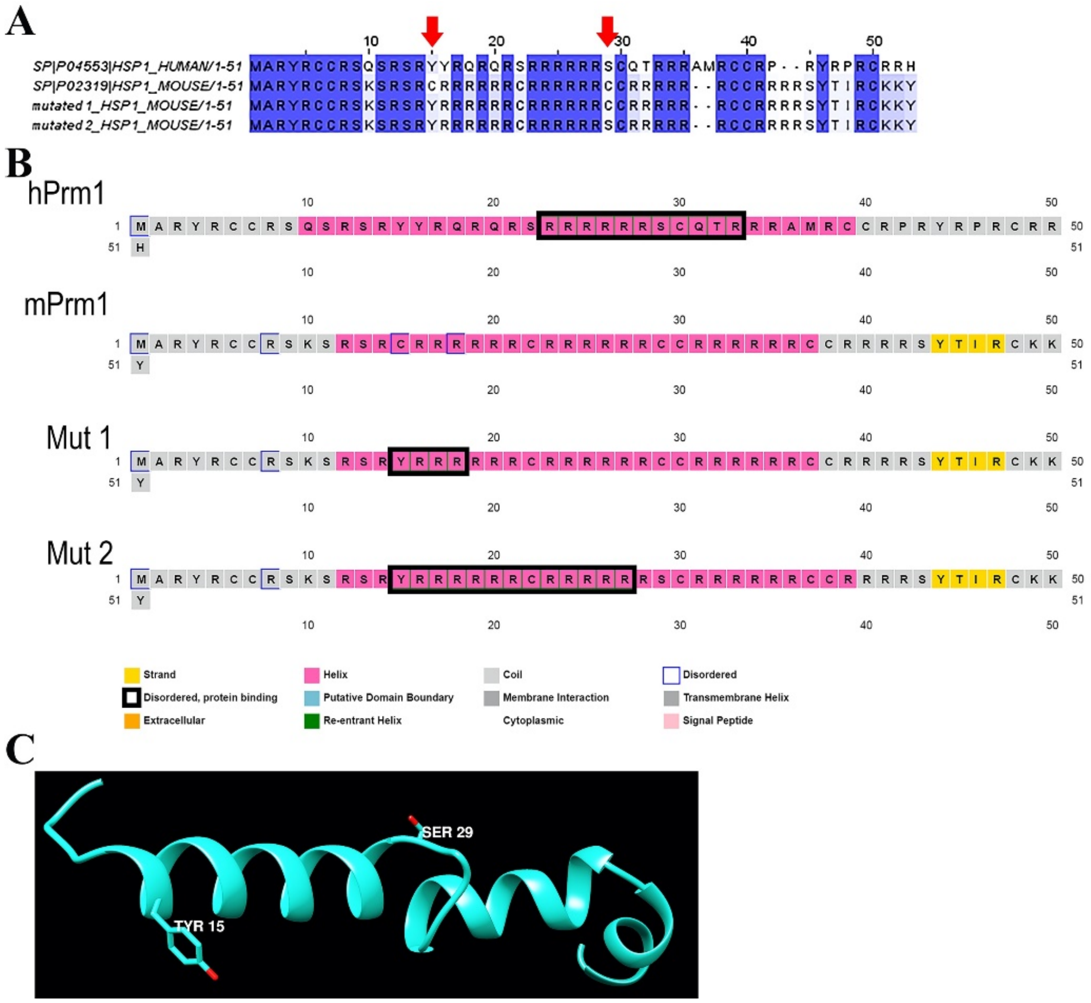


**Supplementary Figure 1.** Protamine 1 and mutated expression vector sequences. (A) Multiple alignment of the wild-type human and mouse protamine 1 sequences and the constructs Mut1-*Prm1* (Mut1) and Mut2-*Prm1* (Mut2). Red arrows indicate cysteine residues (C15 and C29) of the mouse sequence that, in the mutant sequences, are substituted by a tyrosine (C15Y) and a serine (C29S) to resemble the human sequence. (B) Secondary structure prediction according to PSIPRED (http://bioinf.cs.ucl.ac.uk/psipred_new/). Box fillings: grey, coil; pink, helix; yellow, strand. Borders: blue, disordered; black, disordered-protein binding. (C) Homology modeling structure of human protamine 1. The two residues corresponding to the two mutated cysteines in the mouse sequence (Tyr 15 and Ser 29) are shown as ball and stick.

**Supplementary Table 1.** Point mutations in plasmid sequences. Bold characters in the primers highlight the point mutations.

| Plasmid | Mutation | Primers |
|---|---|---|
| Mut1-*Prm1* | g44a c45t | 5'GCAGCAAAAGCAGGAGCAGAT**AT**CGCCGTCGCAG 3' |
| Mut2-*Prm2* | t85a | 5' CGGAGGAGGCGA**A**GCTGCCGGCG 3' |

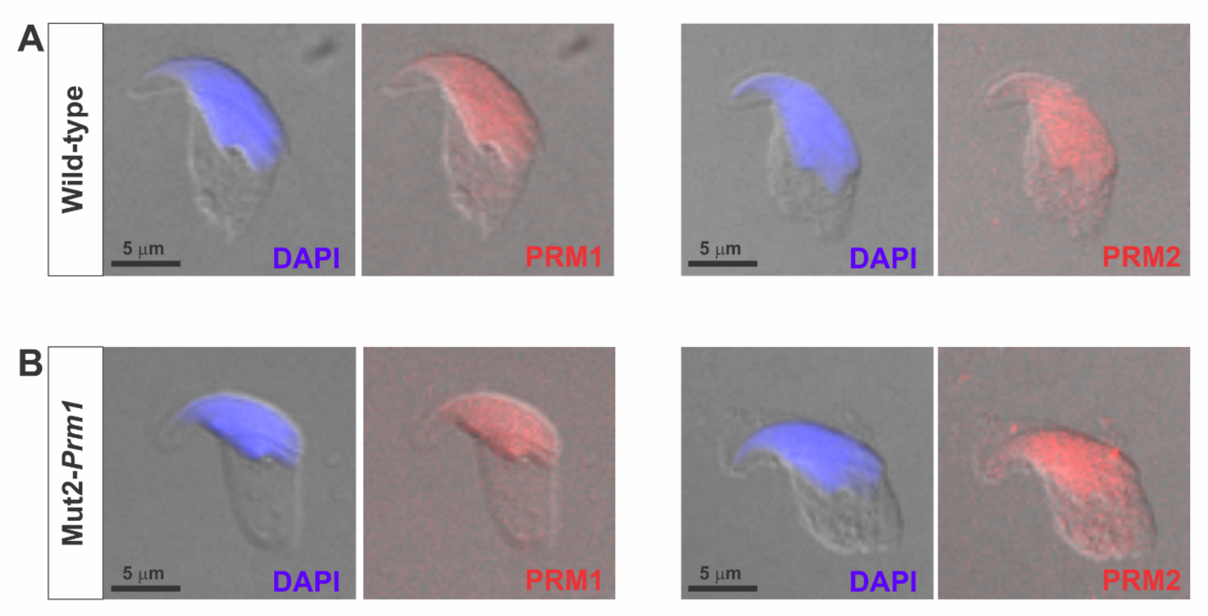


**Supplementary Figure 2.** Mutation of PRM1 in cysteine residues (C15 and C29) does not alter nuclear translocation of PRMs. (A) Representative merge images showing immunofluorescence detection of PRM1 and PRM2 in wild-type spermatids at steps 13-14. (B) Representative merge images showing immunofluorescence detection of PRM1 and PRM2 in Mut2-*Prm1* spermatids at steps 13-14.

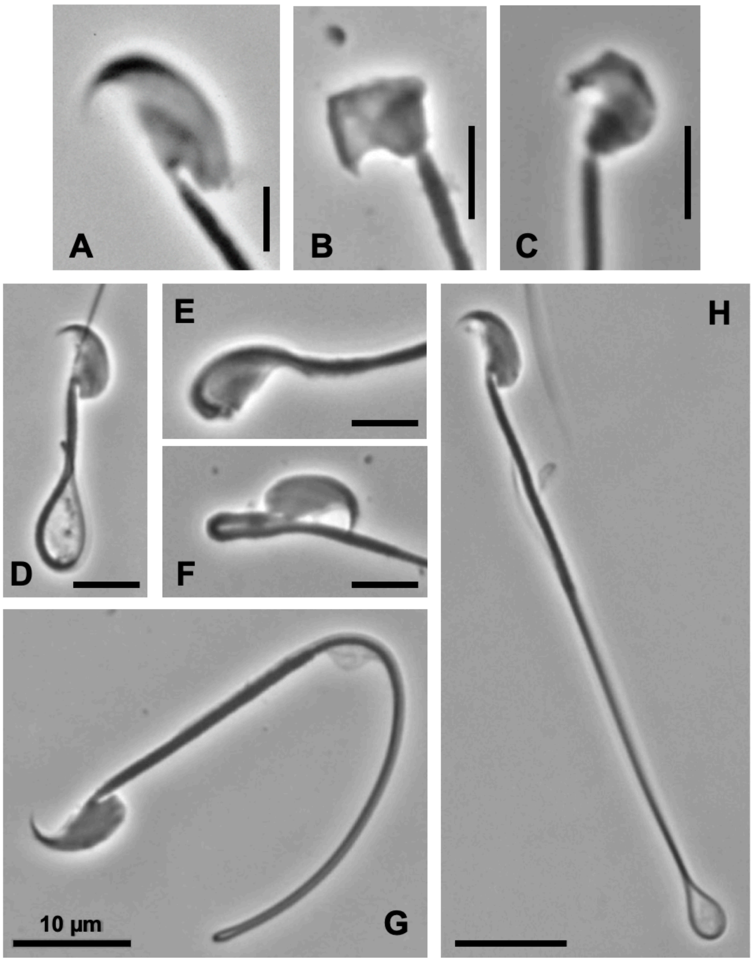


**Supplementary Figure 3.** Abnormalities in mouse spermatozoa. (A) normal sperm head; (B,C) abnormal sperm heads; (D-F) abnormal mipdieces: (G,H) abnormal principal pieces. Scale bars: A-C, 5 µm; D-H,10 µm.

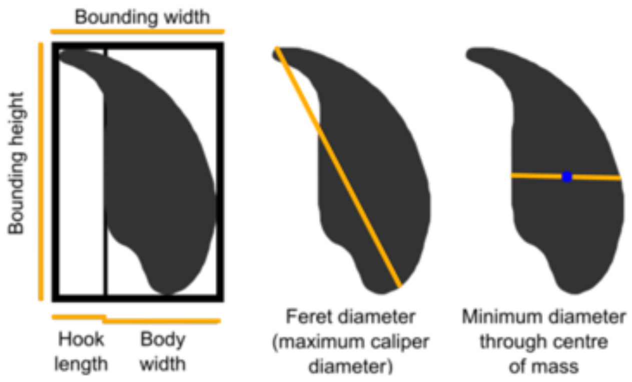


Supplementary Figure 4. Parameters measured by the Nuclear Morphology Analysis software (Skinner et al., 2019).

Supplementary Table 2**.** Description of parameters measured by the Nuclear Morphology Analysis (NMA) software (Skinner et al., 2019).

| **Parameter** | **Description** |
|---|---|
| Area | A; the two-dimensional area of the nucleus |
| Perimeter | P; the length of the nuclear perimeter |
| Max feret diameter | the maximum caliper diameter across the nucleus |
| Min diameter | the shortest caliper diameter through the center of mass of the nucleus |
| Bounding height | H; the height of the bounding rectangle of the vertically oriented nucleus |
| Bounding width | W; the width of the bounding rectangle of the vertically oriented nucleus |
| Width of body | the distance from the vertical region to the x-edge of the bounding rectangle on the body side |
| Length of hook | the distance from the vertical alignment region to the x-edge of the bounding rectangle on the hook side |
| Circularity | $4\pi A/P^2$; the closeness of the nucleus to a circle, between 0 and 1, where 1 is a perfect circle. |
| Ellipticity | H/W; the height (H) divided by width (W) of the nuclear bounding box when the nucleus is vertically oriented |
| Aspect ratio | W/H; the inverse of ellipticity |
| Elongation | (H-W)/(H+W); the bounding height minus the bounding width, divided by the bounding height plus the bounding width |
| Regularity | $(\pi HW)/(4A)$; A measure of how regular the shape is; does it have rotational symmetry |
